# Adaptation and inhibition control pathological synchronization in a model of focal epileptic seizure

**DOI:** 10.1101/312561

**Authors:** Anatoly Buchin, Cliff C. Kerr, Gilles Huberfeld, Richard Miles, Boris Gutkin

**Author notes:** current affiliation: Allen Institute for Brain Science, Modelling Analysis and Theory (United States, Seattle), 615 Westlake Ave N, Seattle, WA 98109.

## Abstract

Pharmacoresistant epilepsy is a common neurological disorder in which increased neuronal intrinsic excitability and synaptic excitation lead to pathologically synchronous behavior in the brain. In the majority of experimental and theoretical epilepsy models, epilepsy is associated with reduced inhibition in the pathological neural circuits, yet effects of intrinsic excitability are usually not explicitly analyzed. Here we present a novel neural mass model that includes intrinsic excitability in the form of spike-frequency adaptation in the excitatory population. We validated our model using local field potential data recorded from human hippocampal/subicular slices. We found that synaptic conductances and slow adaptation in the excitatory population both play essential roles for generating seizures and pre-ictal oscillations. Using bifurcation analysis, we found that transitions towards seizure and back to the resting state take place via Andronov-Hopf bifurcations. These simulations therefore suggest that single neuron adaptation as well as synaptic inhibition are responsible for orchestrating seizure dynamics and transition towards the epileptic state.

**Significance statement:** Epileptic seizures are commonly thought to arise from a pathology of inhibition in the brain circuits. Theoretical models aiming to explain epileptic oscillations usually describe the neural activity solely in terms of inhibition and excitation. Single neuron adaptation properties are usually assumed to have only a limited contribution to seizure dynamics. To explore this issue, we developed a novel neural mass model with adaption in the excitatory population. By including adaptation and intrinsic excitability together with inhibition in this model, we were able to account for several experimentally observed properties of seizures, resting state dynamics, and pre-ictal oscillations, leading to improved understanding of epileptic seizures.

## Introduction

Epilepsy is the fourth most common neurological disorder, and is responsible for a greater total global burden of disease than any neurological conditions except for stroke and migraine (Chin et al., 2014; Beghi et al., 2005; Rothstein et al., 2005). Epileptic seizures are characterized by the increased excitability/excitation in the brain’s recurrently coupled neuronal networks (Lytton, 2009). Typically, experimental seizure models assume that seizures occur due to decreased inhibition (Karnup et al., 1999; Sivakumaran et al., 2015) or increased excitation in the neural networks (Ursino et al., 2006; Hall et al., 2013).

There is also evidence that interneurons increase their firing at seizure initiation (Lillis et al., 2012) and are active during the time course of the epileptic activity (Ziburkus et al., 2006), suggesting that the activity of interneurons contributes importantly to aspects of seizure dynamics. The activity-dependent interplay between the pyramidal cells and interneurons could play an essential role for seizure generation mechanisms (Buchin et al., 2016; Krishnan et al., 2011; Naze et al., 2015). In neural mass models, neuron populations are often treated as rate units lacking intrinsic adaptation (Touboul et al., 2011). The dynamic behavior of the neural populations is determined by the balance between excitation and inhibition. Despite the simplicity of these models, they can be successfully used to reproduce resting and interictal states as well as ictal discharges by producing time series comparable with macroscopic measurements such as electroencephalogram signals (Demont-Guignard et al., 2009).

However, not all types of epileptic seizures can be explained by looking only at the balance between excitation and inhibition (Traub et al., 2005); intrinsic excitability changes on the single-neuron level also play an important role (Krishnan et al., 2011). Studies on human subiculum tissue showed that the complete blockade of type A GABAergic neurotransmission (and thus inactivation of the effects of inhibitory population) precludes seizure emergence while, if applied after seizure initiation, it abolishes rather than enhances the seizure activity. These manipulations usually bring back the neural network in the slice towards pre-ictal events, which have substantially different frequency content than seizure activity (Huberfeld et al., 2011), and which in this case fail to trigger ictal events. In human epileptic tissues, including peri-tumoral neocortex (Pallud et al., 2014), interictal discharges are generated spontaneously. These events are triggered by interneurons which depolarize pyramidal cells with impaired chloride regulation, leading to depolarizing effects of GABA. Once activated, pyramidal cells excite other cells via AMPA-mediated glutamatergic transmission. In these tissues, seizures can be produced by increasing local excitability using modified bathing media. The transition to seizures is characterized by the emergence of specific pre-ictal events initiated by pyramidal cells which synchronize local neurons by AMPA signals. These pre-ictal events cluster before seizure initiation which requires functional AMPA, NMDA as well as GABA_A_ signals. The conventional neural mass models are unable to explain these pre-ictal oscillations because they require the excitatory population to generate periodic oscillations in the absence of inhibition.

The second motivation for incorporating intrinsic excitability into neural mass models is that in epileptogenic areas, such as human subiculum, there is a substantial proportion of neurons with non-trivial intrinsic properties such as spike-frequency adaptation (Huberfeld et al., 2007; Jensen et al., 1994). To take these properties into account, neural mass models need to be enriched by the addition of components such as slow potassium currents (Pinsky et al., 1994).

In addition, seizures are typically accompanied by high potassium concentrations (Xiong et al., 1999; Fröhlich et al., 2006; Florence et al., 2009; Dietzel et al., 1986), which in turn activate calcium currents (Fröhlich et al., 2008; Bazhenov et al., 2004), which in turn affect spike-frequency adaptation and intrinsic bursting. These properties are likely to modulate the single neuron firing and thus further influence the neuronal dynamics. These findings motivate the development of neural mass models that can capture the intrinsic excitability in coupled neural populations.

In this work, we developed a novel neural mass model consisting of an inhibitory neural population and an adaptive excitatory neuronal population (Buchin et al., 2010). We calibrated the parameters of the model to local field potential (LFP) data recorded in human subiculum slices during rest, seizure, and full disinhibition in per-ictal condition. We then analyzed the model as calibrated to each of these three regimes. Our results emphasize the role of intrinsic excitability such as adaptation in the excitatory population, which help explain the transitions between rest, seizure, and full disinhibition states.

## Materials and Methods

### Epileptic tissue

Temporal lobe tissue blocks containing the hippocampus, subiculum, and part of the entorhinal cortex were obtained from 45 people of both sexes with pharmacoresistant medial temporal lobe epilepsies associated with hippocampal sclerosis (age, 18–52 years; seizures for 3–35 years) undergoing resection of the amygdala, the hippocampus, and the anterior parahippocampal gyrus. All of the individuals gave their written informed consent and the study was approved by the Comité Consultatif National d’Ethique.

### Tissue preparation

The post-surgical tissue was transported in a cold, oxygenated solution containing 248 mM d-sucrose, 26 mM NaHCO_3_, 1 mM KCl, 1 mM CaCl_2_, 10 mM MgCl_2_ and 10 mM d-glucose, equilibrated with 5% CO_2_ in 95% O_2_. Hippocampal-subicular-entorhinal cortical slices or isolated subicular slices (400 μm thickness, 3×12 mm length and width) were cut with a vibratome (HM650 V, Microm). They were maintained at 37 °C, and equilibrated with 5% CO_2_ in 95% O_2_ in an interface chamber perfused with a solution containing 124 mM NaCl, 26 mM NaHCO_3_, 4 mM KCl, 2 mM MgCl_2_, 2 mM CaCl_2_ and 10 mM d-glucose. Bicuculline or picrotoxin was used to block GABA_A_ receptors. Ictal-like activity was induced by increasing the external K^+^ concentration to 8 mM and reducing the Mg^2+^ concentration to 0.25 mM to increase the cellular excitability (similar to Huberfeld et al., 2011).

### Recordings

Up to four tungsten electrodes etched to a tip diameter of ^~^5 μm were used for the extracellular recordings. The signals were amplified 1,000-fold and filtered to pass frequencies of 0.1 Hz to 10 kHz (AM systems, 1700). The extracellular signals were digitized at 10 kHz with a 12–bit, 16-channel A-D converter (Digidata 1200A, Axon Instruments), and monitored and saved to a PC with Axoscope (Axon Instruments).

### Data analysis

Records were analyzed using pCLAMP 10 software and scripts written in Matlab 2016a. Power spectrum estimation was performed using fast Fourier transforms. The major frequencies of oscillations were computed via the multitaper method (Thomson 1982).

### Simulations and analysis

Neural population model simulations were performed in XPPAUT 8.0 using the direct Euler method of integration, with a time step of 0.05 ms. Smaller time steps were tested and provided substantially similar results. Bifurcation analysis was performed in the AUTO package (http://www.math.pitt.edu/~bard/xpp/xpp.html). In all simulations the initial conditions were systematically varied to check stability of numerical results. The model code is available on GitHub (https://github.com/abuchin/EI-with-adaptation). The data for the model was taken from one representative patient in the brain slice demonstrating resting state, seizure and pre-ictal oscillations.

### Neural mass model

In the model we considered interacting excitatory and inhibitory neural populations coupled by AMPA and GABA_A_ synapses. All model variables and parameters are presented in Tables 1 and 2. Each population was characterized by the average membrane potential of a population of leaky integrate-and-fire (LIF) neurons, similar to (Touboul et al., 2011, Chizhov et al., 2007) with approximations for adaptive currents taken from (Buchin et al., 2010):

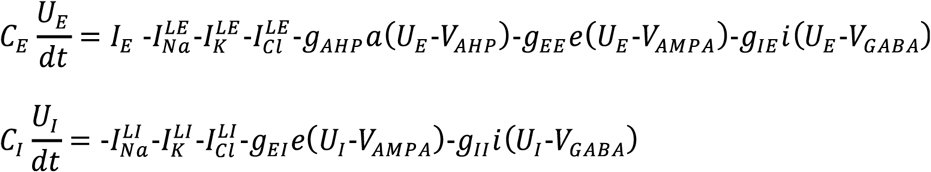

where

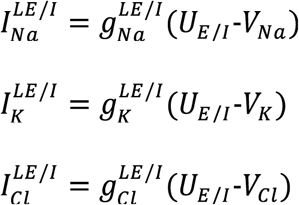

**Table 1.**
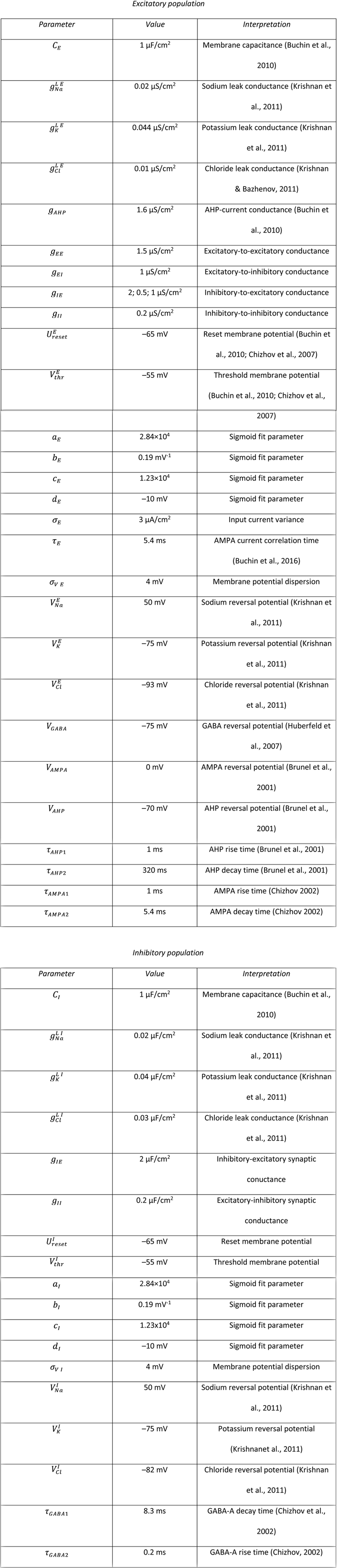
Population model parameters

**Table 2.** Population model variables

| *Variable* | *Interpretation* |
| --- | --- |
| $U_E$, mV | Average membrane potential of the excitatory population |
| $U_I$, mV | Average membrane potential of the inhibitory population |
| $e$ | Excitatory population synaptic gating variable |
| $i$ | Inhibitory population synaptic gating variable |
| $a$ | Excitatory population adaptation gating variable |
| $I_E(t)$, $\mu\text{A}/\text{cm}^2$ | Random excitatory input |
| $\nu_E(t)$, Hz | Firing-rate of the excitatory population |
| $\nu_I(t)$, Hz | Firing-rate of the inhibitory population |

The firing rate of each population is computed based on the interspike interval distribution of the neural population (Gerstner et al., 2002):

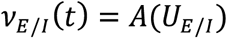

where

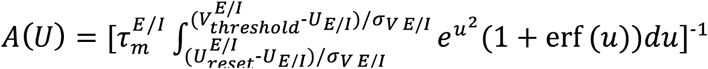

and

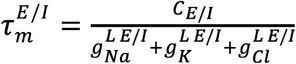

In all simulations *ν_E/I_*(*t*) has been approximated by the following sigmoid function:

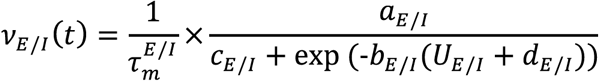

The population firing rate determines the adaptive (a), excitatory (e) and inhibitory (i) conductances. Their dynamics are computed using the second-order approximation (Wendling et al., 2002; Chizhov, 2013):

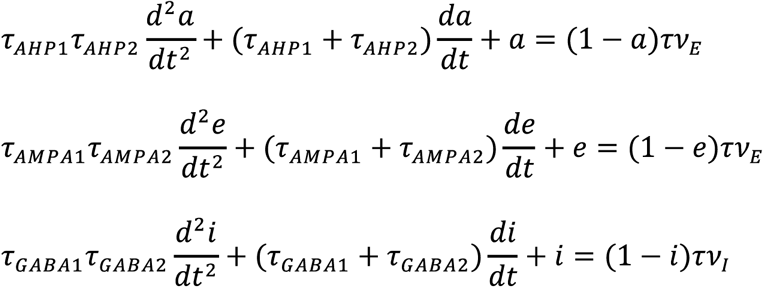

To mimic the afferent excitatory input, the excitatory population also received stochastic excitatory input modeled as an Ornstein-Uhlenbeck process (Buchin et al., 2010):

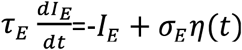

To mimic elevated extracellular potassium from epileptogenic slice experiments, in the population model, we increased potassium reversal potential in both populations 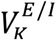 from –90 mV to –75 mV, i.e. from *K_o_*=4 mM to *K_o_*=8 mM. This value of 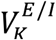 was computed based on Nernst equation, 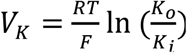, where 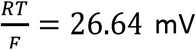 and *K_i_* = 138 mM (Krishnan et al, 2011).

All model parameter values and variable names are present in Table 1 and 2. The initial parameter set was chosen manually to reproduce the pre-ictal like oscillations due to balance between *g*_*EE*_ and *g*_*AHP*_, seizure and resting state were fit such that *g*_*EI*_ parameter variations would make a transition between seizure and resting state.

### Local field potential model

The LFP was calculated based on the activity of the excitatory population. We assumed that pyramidal cells activity dominates the extracellular field (Buzsáki et al., 2012). The dominant theory is that the LFP component is dominated by the single neuron dipole contribution (Buzsáki et al., 2012). Since the neural mass model averages over single neurons, the dipole moment cannot be directly modeled. Thus, to approximate the LFP being recorded near somas of the excitatory populations, we used the assumption that the average membrane potential of the excitatory population is proportional to the LFP, i.e. LGP ∝ *U_E_* (Ursino et al., 2006; Demont-Guignard et al., 2009; Wendling et al., 2012; Ratnadurai-Giridharan et al., 2014).

## Results

### Construction of the population model

We developed а model of interacting excitatory and inhibitory population inspired by Wilson-Cowan approach (Wilson et al., 1972). It consists of excitatory and inhibitory populations coupled by synaptic connections, as shown in Fig. 1A. The firing rate in each population depends on the average membrane potential *U_E/I_*, which is governed by the subthreshold dynamics of leaky integrate-and-fire (LIF) neuron population similar to (Gerstner et al., 2002; Chizhov, 2013), as explained in the Materials and Methods. Firing rates of the excitatory and inhibitory populations are determined using the values of *U_E/I_* put through function *A*(*U_E_*/*I*) (Gerstner et al., 2002; Johannesma, 1968). To make the model numerically stable and amenable to bifurcation analysis we used a sigmoid function to estimate the population firing rate provided by the *A*(*U_E_*_/*I*_) approximation. To justify the choice of sigmoid parameters we used least-squares to match it with the analytical solution (Johannesma, 1968), as shown in Fig. 1C, D. The sigmoid approximation allows one to efficiently take into account zero and linear parts of the potential-to-rate transfer functions *υ_E_*_/*I*_, and provides saturation due to the single neuron refractory period (Renart et al., 2004). The sigmoid functions of excitatory and inhibitory populations are shown in Fig. 1C and 1D. The difference between the excitatory and inhibitory populations was taken into account by adjusting passive conductances for sodium, potassium, and chloride leak currents estimated in (Krishnan et al., 2011) based on dynamic ion concentration model.

The subthreshold *U_E_*_/*I*_ dynamics determine the synaptic *g_EE_e*(*υ_E_*), *g_EE_e*(*υ_E_*_/*I*_), *g_II_i*(*υ_I_*), *g_IE_i*(*υ_I_*) and intrinsic *g_AHP_e*(*υ_E_*_/*I*_) conductances, shown in Fig. 1A, computed according to the population firing-rates *υ_E_*_/*I*_. Similar to spiking neural network models (Bazhenov et al., 2004), (Ratnadurai-Giridharan et al., 2014), adaption in our population model reduces neural firing in the excitatory population after periods of activity. Excitatory population receives external random synaptic input to model excitation from the rest of the brain similar to (Touboul et al., 2011; Jansen et al., 1995). To mimic the experimental epileptogenic conditions of human subiculum slice experiments, the potassium reversal potential was elevated from –95 mV to –75 mV both in the excitatory and inhibitory populations to provide excitatory drive to reproduce the experimental conditions. Elevation of extracellular potassium also leads the increase of intracellular chloride reducing the efficiency of inhibition due to elevated GABA_A_ reversal potential (Huberfeld et al., 2007), Buchin et al., 2016). To generate the model output comparable with experimental data, we computed the LFP generated by the excitatory population (Buzsáki et al., 2012). This approximation assumes that all pyramidal cells in the excitatory population contribute equally to the recorded LFP signal, as shown in Fig. 1B. Thus the total LFP near somas depends on the average value of the membrane potential in the excitatory population with a certain dimensionality constant, i.e. *LFP* ∝ *kU_E_*_/*I*_.

**Figure 1.**
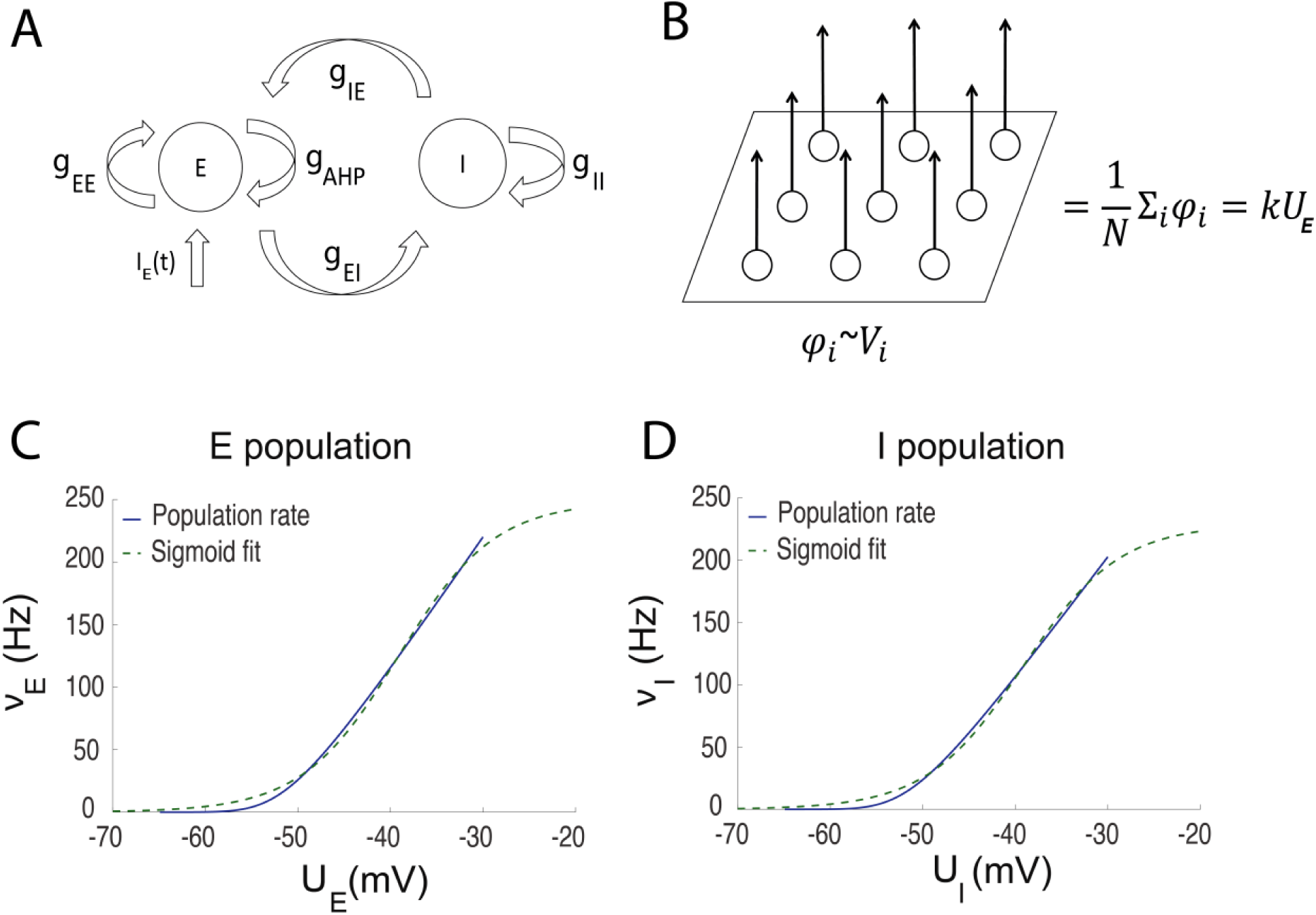
Structure of the population model. (A) Scheme of interacting neural populations. E, I – excitatory and inhibitory populations. *g_EE_*, *g_EI_* – excitatory to excitatory and excitatory to inhibitory maximal conductances. *g_II_*, *g_IE_*– inhibitory-to-inhibitory and inhibitory-to-excitatory maximal conductances. *I_E_*(*t*)– synaptic noise input to the excitatory population. AHP – after hyperpolarization current (Buchin et al., 2010). (B) LFP model:*φ_i_*-contribution of a single excitatory cell, N – the number of neurons, *U_E_* - the average membrane potential in the excitatory population. (C, D) Sigmoid approximation of potential-to-rate function (Johannesma, 1968) of the excitatory (C) and inhibitory population (D).

### Reproduction of epileptic oscillations

When the excitatory and inhibitory synaptic currents were dynamically balanced, the activity stayed in the low-firing regime, as indicated by LFP power spectrum. The recorded pyramidal cell during this period demonstrated sparse firing activity, partially time-locked with the discharges on the LFP. We call this activity in the model the balanced or *resting state*. In this regime the model does not generate epileptic oscillations. To evaluate the model performance in this resting state, we compared the synthetic LFP with the experimental LFP recorded between seizures, as shown in Fig. 2A. Similar to the experimental data, we found that in the resting state, the model generates broadband oscillations, with the highest power in the 1–15 Hz frequency band. In this regime, the average membrane potential of the excitatory population *U_E_*_/*I*_ stays in the range from –60 to –50 mV.

**Figure 2.**
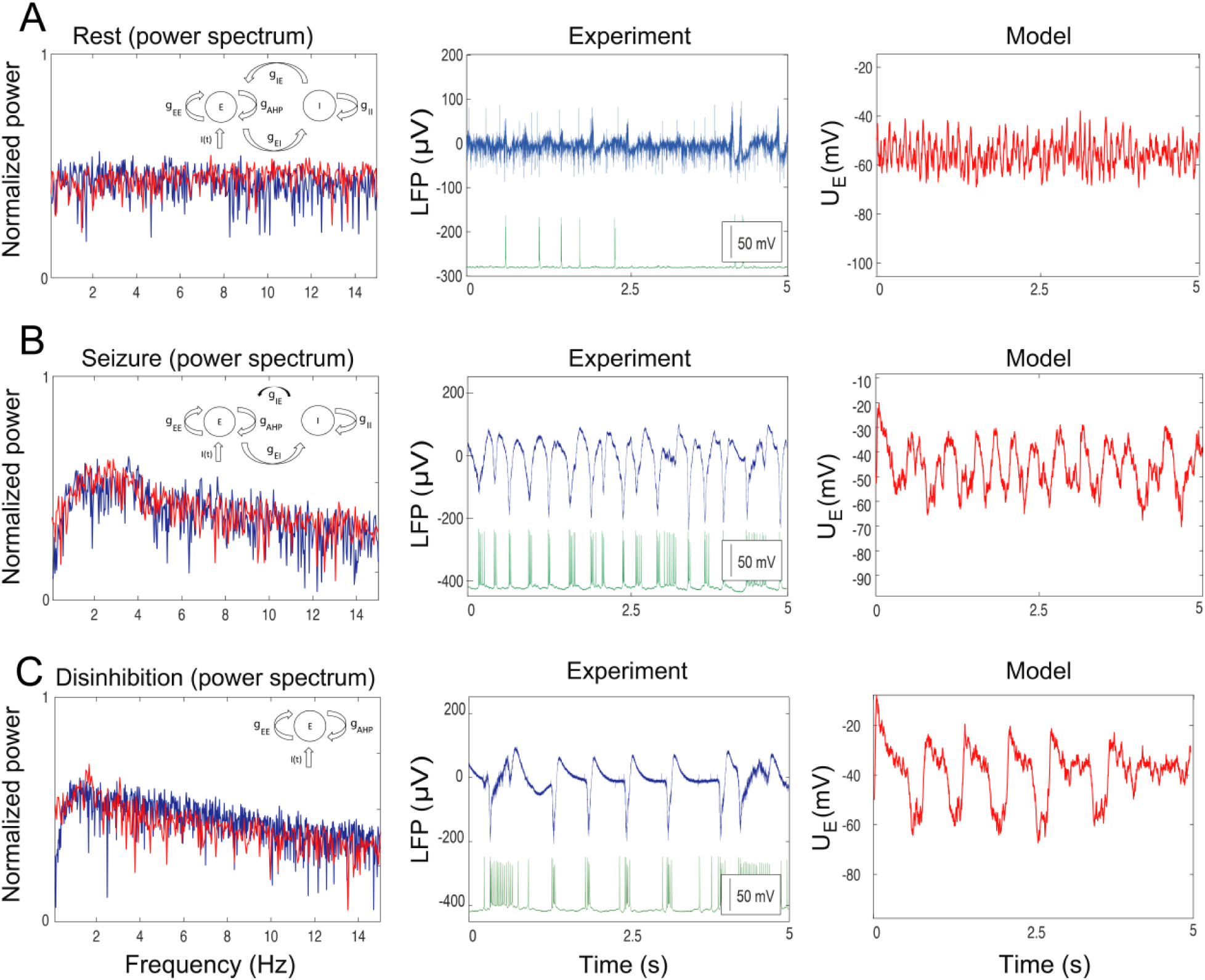
Neural mass model in various excitatory regimes. (A) Activity of a neural population in the resting state. (B) Seizure state. (C) Disinhibited state. LFP is present together with intracellular recording from the pyramidal cell. Each plot contains the model scheme, power spectrum and time traces provided by the excitatory population *U_E_* as well as experimental LFP. Red traces correspond to the model, blue traces to the experiment. Green traces are the intracellular recordings from the pyramidal cells. Corresponding model parameters (A): *g_EE_* =1.5 uS/cm^2^; *g_EI_* =1 uS/cm^2^; *g_IE_* =2 uS/cm^2^; *g_II_* =0.2 uS/cm^2^; *g_AHP_* =1.6 uS/cm^2^; (B): *g_EE_* =1.5 uS/cm^2^; *g_EI_* =1 uS/cm^2^; *g_IE_* =0.5 uS/cm^2^; *g_II_* =0.2 uS/cm^2^; *g_AHP_* =1.6 uS/cm^2^; (C): *g_EE_* =1.5 uS/cm^2^; *g_EI_* =1 uS/cm^2^; *g_IE_* =0 uS/cm^2^; *g_II_* =0.2 uS/cm^2^; *g_AHP_* =1.6 uS/cm^2^.

We found that the model was not capable of generating interictal discharges using this parameter set. It has been recently suggested that interneurons play the key role in generating interictal activity (Cohen et al., 2002; Huberfeld et al., 2011). In the presence of GABA_A_ blockade these events were completely blocked, indicating that they depend on combination of GABAergic and glutamatergic signaling. In the recent population model (Chizhov at al. 2017) it was proposed that interictal discharges could be initiated by the inhibitory population, thus explaining interneuron firing prior to pyramidal cell firing (Huberfeld et al., 2011). In our model we have not explored this scenario, i.e. when the inhibitory population is also receiving the background synaptic input. These mechanisms would likely play an important role for seizure initiation; however, incorporating all mechanisms at once would make the model impossible to study analytically. Therefore, we have not considered interictal discharges prior to seizure, while aiming to specifically describe other types of oscillations.

To reproduce the seizure state in the model, we reduced the synaptic inhibition of the excitatory population by decreasing the synaptic conductance parameter *g_IE_*, as shown in Fig. 2B (black arrow). All other parameters of the model remained the same. In this case the model moved into an oscillatory regime in which the power spectrum of the oscillations changed dramatically to include strong oscillations in the 1–4 Hz frequency band, which is typical for ictal discharges (Huberfeld et al., 2011).

We compared the model power spectrum with the measured LFP recorded during the initial phase of the ictal discharge with the hypersynchronous activity onset. During this activity regime the recorded pyramidal cells generated strong bursts of spikes temporally locked to the LFP, as shown in Fig. 2B. The population model displayed discharges with the same frequency band as in the LFP, indicating large amount of synchrony in the excitatory population (Buzsáki et al., 2012). Note that we considered only the initial phase of the seizure; the whole ictal event is shown in Fig. 3E.

**Figure 3.**
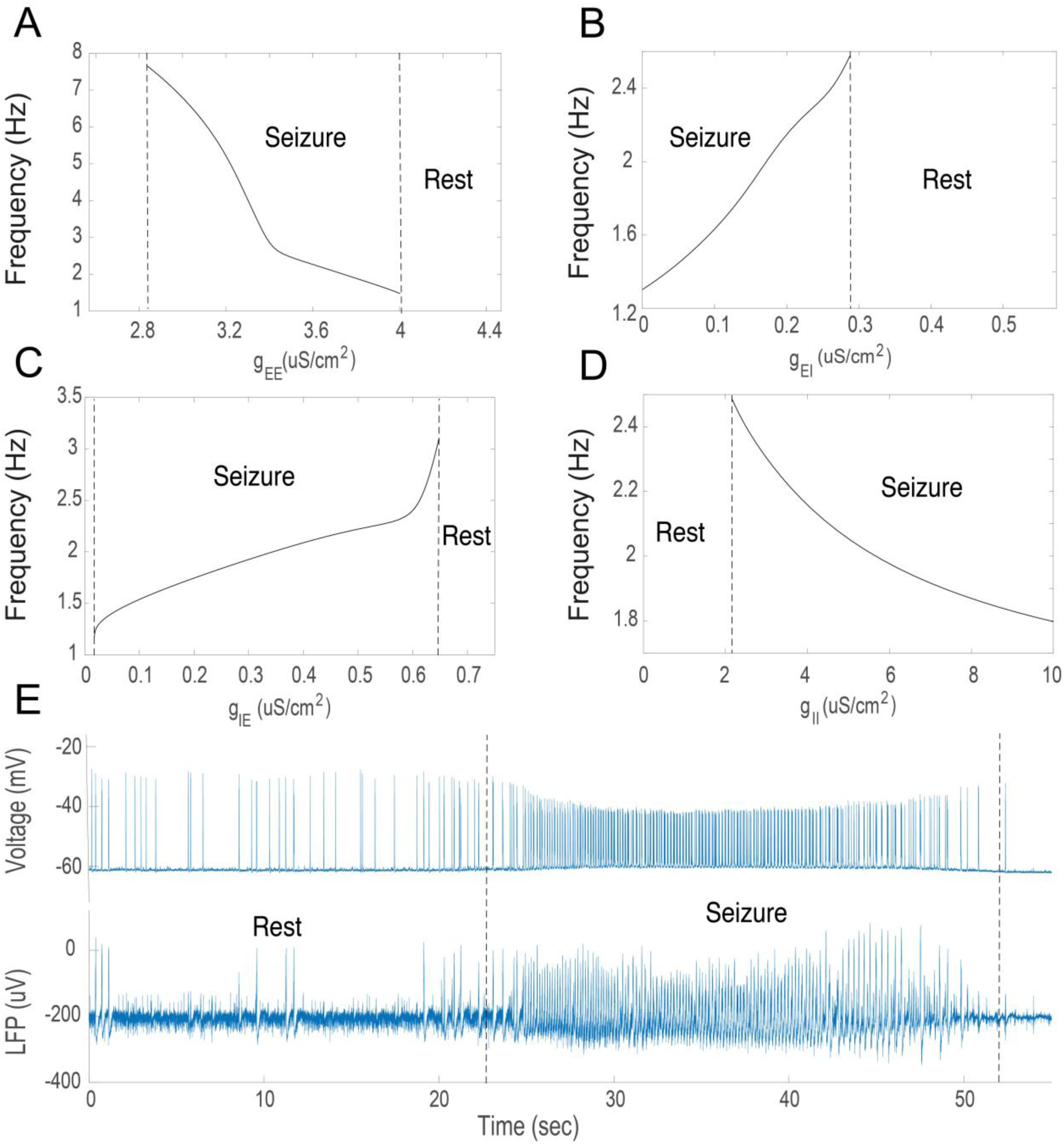
Oscillatory frequencies of the population model. (A–D) Oscillatory frequencies of the population model in the absence of the synaptic noise (*I_E_*(t)=0) as a function of the synaptic gain, *g_EE_, g_EI_, g_IE_, g_II_*. (E) Simultaneous intracellular recording from the pyramidal cell and LFP during transition between the resting states and seizure, marked by dotted lines.

To further test the validity of our model, we explored its dynamics with inhibitory activity completely blocked, as shown in Fig. 2C. In these simulations the initial conditions and parameter values of the model were set to the seizure state, but with the conductance *g_IE_* (from the inhibitory to the excitatory population) set to zero to mimic the experimental conditions. In this case the GABAergic effects of the inhibitory population in the slice has been fully blocked by bicuculine after seizures have been previously established (Huberfeld et al., 2011). In response to this change, the activity in the slice became highly synchronized and reduced to regular pre-ictal discharges. During these oscillations the pyramidal cells generated large bursts of activity, temporally coupled with the LFP, Fig. 2C. In the model, similarly to the experimental preparation, the blockade of the GABAergic signaling mimicked by the abolition of the inhibitory population led to the development of a slow oscillatory rhythm with a peak frequency around 1 Hz. These events have been previously reported as pre-ictal discharges (Huberfeld et al., 2011). This rhythm has much slower frequency than seizures, and is usually within 1–4 Hz frequency range (Buchin et al., 2016; Huberfeld et al., 2011). In addition, these events recur regularly for long periods with very limited modulation.

We call this regime of activity pre-ictal discharges because similar activity takes place before transition towards an ictal state (Huberfeld et al., 2011). In this regime, the dynamics of the excitatory population are determined only by the balance between self-excitation, *g_EE_e*(*ν_E_*_/*I*_), after-hyperpolarization current (AHP) (Buchin et al., 2010; Chizhov et al., 2008), *g_AHP_ e*(*ν_E_*_/*I*_), and the afferent synaptic current *I_E_*(*t*). Hence these pre-ictal oscillations in the model are driven by the synaptic noise and adaptation. The excitatory input to the excitatory population *I_E_*(*t*) drives the upswings of *U_E_* due to recurrent excitatory synapses, with activity then being terminated by AHP currents. These transitions take place randomly due to stochastic nature of the synaptic input.

For quantitative comparisons between the model and experiment we used the linear fit to the power spectrum over frequencies and peak estimation, as shown in Table 3. We found that there is substantial intersection between linear fits applied to the power spectrums in resting, seizure, and pre-ictal states, as shown in Fig. 2. We found that there is a substantial overlap between these frequencies, providing validation for the model. Note that we compared the overall spectral characteristics between the model and experiment by variation of only one parameter, *g_IE_* to reproduce transitions between the pre-ictal, resting and seizure states. If more parameters are varied at the same time, it would be possible to get a better match between the model and experiment.

**Table 3.** Power spectrum analysis

|  | Model, peak<br>amplitude, Hz | Experiment, peak<br>amplitude, Hz | Model,<br>spectrum<br>linear fit, 1/Hz | Experiment,<br>spectrum<br>linear fit, 1/Hz |
| --- | --- | --- | --- | --- |
| Rest | - | - | -0.005; -0.002 | -0.005; -0.002 |
| Seizure | 3.01; 3.52 | 2.95; 3.75 | -0.005; -0.002 | -0.003; -0.002 |
| Pre-ictal state | 1.33; 1.43 | 1.21; 1.79 | -0.007; -0.003 | -0.01; -0.008 |

Overall oscillations in our population model are controlled by the balance between synaptic currents, adaptation and external synaptic input. When synaptic and intrinsic conductances are balanced, the population demonstrates resting state activity, characterized by a flat power spectrum. When there is an imbalance between excitation and inhibition, populations start developing oscillatory rhythms associated with ictal discharges with a frequency of 3-4 Hz. However, complete loss of inhibition leads to the development of another population rhythm, pre-ictal discharges with 1 Hz frequency, controlled by adaptation and recurrent excitation. Thus the dynamic state of a neural population depends on the interplay between the intrinsic and synaptic excitability within populations as well as external synaptic input.

### Analysis of the population model

In order to delineate the mechanisms giving rise to the different oscillatory modes in the model, we used continuation techniques and bifurcation analysis. Since it is impossible to use the standard techniques to identify bifurcations in the presence of noise, we analyzed the model in the absence of an external input *I_E_*(*t*). This allowed us to compute the model behavior in the stationary regime and characterize bifurcations happening during transitions between different oscillatory regimes. The initial parameters were chosen to correspond to the resting state. The parameter variations were calculated around this point in the parameter space for *g_EE_*, *g_EI_*, *g_IE_* and *g_II_* bifurcation diagrams, with other parameters held fixed. Analysis of *g_AHP_* and *V*_*GABA*_ variations was implemented for another parameter set, where *g_IE_* = 0.5 uS/cm^2^ and *g_IE_* = 1uS/cm^2^; other parameters remained the same.

The frequency of seizure oscillations depends on the strength of the synaptic currents in the population model. There is a nonlinear relationship between seizure major frequency and the recurrent excitatory conductance *g_EE_*, as shown in Fig. 3A. When the *g_EE_* is increased up to 2.8 mS/cm^2^, the model responds with an oscillatory frequency near 7.5 Hz. When self-excitation is further increased up to 4 mS/cm^2^, seizure-related oscillations disappear since the system moves to the high activity state due to sigmoidal saturation of the transfer function, as shown in Fig. 1C and 1D. The amount of stimulation of the inhibitory population also influences the oscillatory frequency. When *g_EI_* is in the range from 0 to 0.29 mS/cm^2^, shown in Fig. 3B, the population model generates seizure activity with frequencies from 1.2 to 2.5 Hz. Note that seizure oscillations are possible even when *g_EI_* = mS/cm^2^.

Inhibitory synaptic connections also affect the oscillatory frequency of seizure activity. When *g_IE_* is as low as ^~^0.6 mS/cm^2^, as shown in Fig. 3C, the seizure activity starts around 3 Hz; it decreases to approximately 1 Hz when *g_EI_* is close to zero (when *g_IE_* = 0 mS/cm2, there is no seizure activity in the model). The amount of recurrent inhibition also determines the seizure oscillatory frequency, as shown in Fig. 3D. Seizure activity can be initiated by sufficient inhibition, i.e. when *g_II_* is near 2 mS/cm^2^, seizures of 2.5 Hz are observed. When *g_II_* increases, the seizure frequency decreases; for example, at 10 mS/cm^2^, seizure activity is approximately 1.8 Hz.

In the previous sections, the population model was calibrated to data for short periods of seizure activity, where the frequency was not substantially changing, as shown in Fig. 2B. Yet, one can see that in the experiment, seizure activity is not stationary and its frequency changes over time. The time course of a typical seizure is shown in Fig. 3E. Before the seizure starts there is a resting state, characterized by occasional interictal (Cohen et al., 2002) and pre-ictal discharges (Huberfeld et al., 2011). When seizure starts at 22 s, it is characterized by fast oscillations of the extracellular field in the range of 5–6 Hz in the initial phase. During the time course of seizure activity, it gradually decreases to 1 Hz frequency, and from 52 s it gradually stops.

To study the amplitude of pathological oscillations, we performed a bifurcation analysis and tracked changes of the average membrane potential in the excitatory population, *U_E_*, as shown in Fig. 4. the self-excitation conductance *g_EE_* (Fig. 4A). We found that increasing *g_EE_* leads to the development of ictal oscillations when its value increases beyond approximately 2.8 mS/cm^2^. During the gradual increase of *g_EE_*, the constant steady state loses stability via the supercritical Hopf bifurcation (Izhikevich, 2007). After passing this point the neural populations start developing seizure oscillations. This activity regime is stable for large *g_EE_* variations, implying that seizure dynamics are possible for a large range of recurrent excitation. When *g_EE_* becomes higher than a critical value (more than 4.1 mS/cm^2^) the seizure state goes to the high activity state with no oscillations. This happens due to the sigmoid approximation of the population rate (Johannesma, 1968), when *ν_E_*_/*I*_ reaches the saturation level, as shown in Fig. 1C, D.

**Figure 4.**
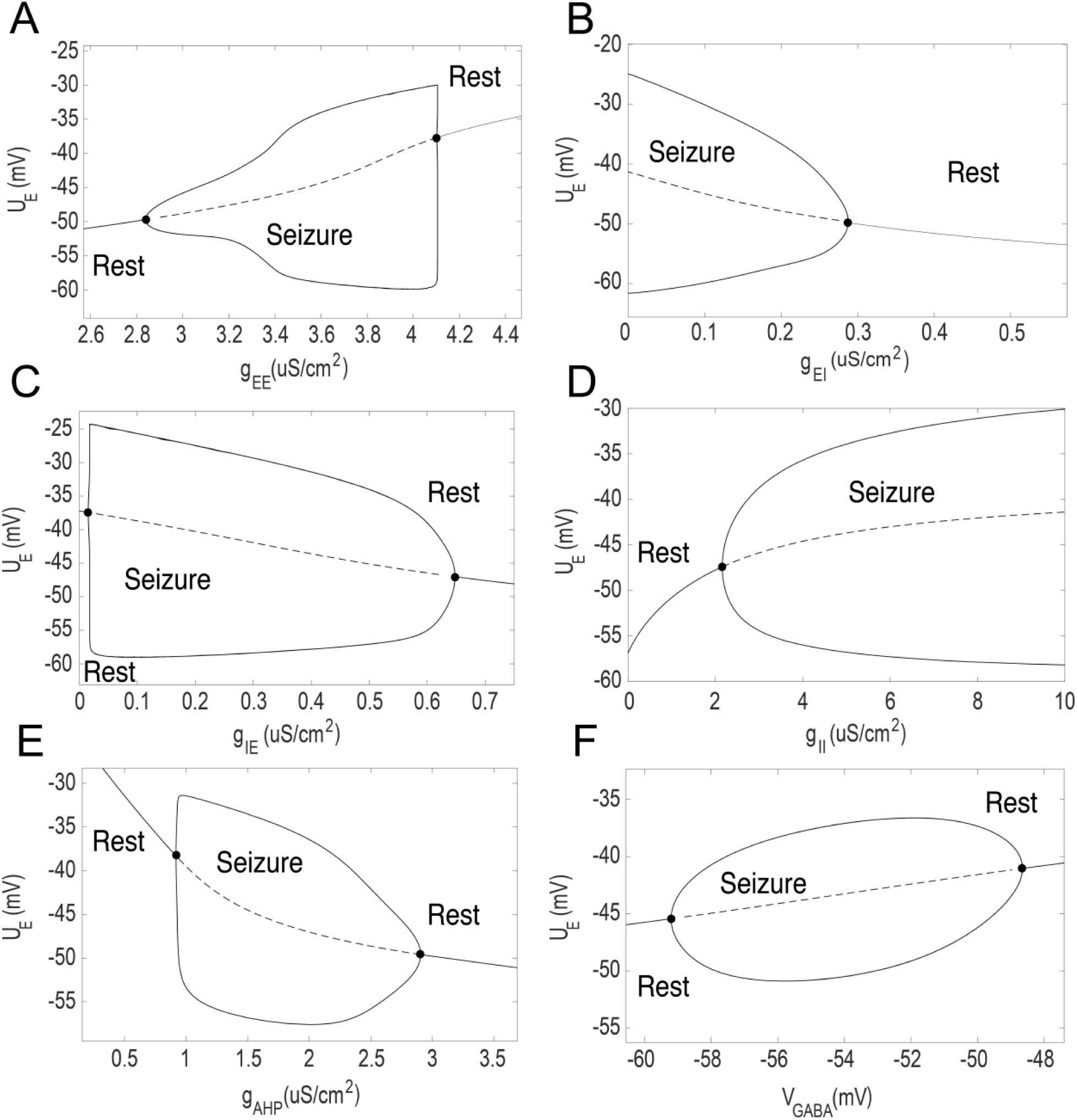
Analysis of the population model. (A–D) Bifurcation diagrams for the variations of the maximal synaptic conductances, including recurrent excitation *g_EE_*, excitation from excitatory to inhibitory population *g_EI_*, inhibition from inhibitory to excitatory population *g_IE_*, and the recurrent inhibition in the inhibitory population *g_II_*, respectively. (E, F) Bifurcation diagrams for adaptation in the excitatory population *g_AHP_* and GABA reversal potential *V_GABA_* from the inhibitory-to-excitatory connection, *g_IE_i*(*U_E_-V_GABA_*). Diagrams A–D were calculated for *g_IE_* =2 *μ*S/cm^2^; E, 0.5 *μ*S/cm^2^; and F, 1 *μ*S/cm^2^. The value of *U_E_* characterizes the average membrane potential in the resting state and maximal/minimal values of *U_E_* during the oscillations. Dots correspond to the Andronov-Hopf bifurcations. Solid and dotted lines depict to the stable and unstable solutions.

Second, we considered the excitatory to inhibitory conductance *g_EI_* (Fig. 4B). In this case, seizure activity is blocked when *g_EI_* is larger than 0.3 mS/cm^2^. If *g_EI_* is smaller than 0.3 mS/cm^2^, it leads to seizure activity via a subcritical Hopf bifurcation. Similar to the *g_EE_* bifurcation diagram, seizure dynamics are possible for a large range of *g_EI_*. These results show that a decrease in the excitatory conductance from excitatory to inhibitory populations is sufficient to provoke seizure activity. Note that even if *g_EI_* = 0 mS/cm^2^, the excitatory population still receives the input from the inhibitory one because potassium reversal potential is elevated. These changes in potassium reversal potential drive both excitatory and inhibitory population even if synaptic drive is not present. For example, when *g_EI_* = 0 mS/cm^2^ the increased potassium reversal potential still drives the inhibitory population, providing the inhibitory input to the excitatory population. It happens because it decreases the leak current thus depolarizing the membrane potential of excitatory and inhibitory neurons. Therefore, seizure oscillations are still present because inhibition is still present. Seizure frequency in this case is near 1.25 Hz (Fig. 3B) and *U_E_* oscillates between –61 and –25 mV.

Third, we considered inhibitory to excitatory conductance *g_IE_* (Fig. 4C). When *g_IE_* = 0, the model shows resting state activity. This corresponds to the condition when the inhibitory population does not have any influence on the excitatory one. Experimentally this scenario is achieved when inhibitory neurotransmission is completely blocked. Therefore, in the complete absence of inhibition, seizure activity could not be generated. In turn, pre-ictal oscillations are not possible without the contribution of the external synaptic noise *I_E_*(t) when *g_IE_* = 0 mS/cm^2^. When there is stochastic synaptic input, it occasionally brings the system to the oscillatory regime associated with seizures (Fig. 2C). Then oscillations are promoted due to recurrent excitation and terminated via AHP adaptation mechanism. Thus without participation of the inhibitory population, the model is incapable of seizure generation. In turn, pre-ictal oscillations do not require inhibition, but strongly depend on the recurrent excitatatory-to-excitatory connections *g_EE_*, adaptation *g_AHP_* and the external synaptic input *I_E_*(t). When inhibitory to excitatory conductance *g_IE_* becomes strong enough, around 0.65 mS/cm^2^, seizure oscillations become truncated and the system moves back to the resting state via subcritical Andronov-Hopf bifurcation.

Fourth, we evaluated the role of recurrent inhibitory conductance *g_II_* for seizure dynamics (Fig. 4D). When there is substantial amount of self-inhibition in the inhibitory population, it leads to an increase of excitation in the whole system because of synaptic coupling. If *g_II_* is above 2.1 mS/cm^2^, it leads to the development of seizure oscillations via a supercritical Hopf bifurcation. Seizure activity in this case persists for the large variations in *g_II_* variations, from 2.2 until more than 10 mS/cm^2^.

We then analyzed the effect of adaptation in the excitatory population. We found the regime in the parameter space of the model for which *g_AHP_* becomes the critical parameter for seizure oscillations. To find this regime we slightly modified the parameter set, where *g_IE_*=0.5 mS/cm^2^ instead of 2 mS/cm^2^, corresponding to the dynamic regime where *g_AHP_* becomes the bifurcation parameter. In this case *g_AHP_* could substantially affect seizure oscillations. When *g_AHP_* is in the range from 1 to 3 mS/cm^2^, there is a large region in the parameter space that produces seizure oscillations. If *g_AHP_* is larger than 3 mS/cm^2^, the seizure dynamics becomes truncated due to the inhibitory effect of adaptation. Yet when adaptation is not strong enough, *g_AHP_* is lower, and the model demonstrates seizure oscillations. If *g_AHP_* is lower than 1 mS/cm^2^, seizure oscillations become impossible and the model stays in the high activity state without oscillations. Additionally, we found that in the complete absence of adaptation, seizure oscillations are still possible in the model (results not shown), but pre-ictal oscillations could not be generated. To be able to reproduce seizure oscillations together with pre-ictal oscillations induced by GABA_A_ blockade, adaptation in the excitatory population is required.

We further studied the critical role of *V_GABA_* for seizure generation. It has been recently found that changes in *V_GABA_* are associated with the rhythm generation in the hippocampus (Huberfeld et al., 2007; Cohen et al., 2002). The analysis was performed for slightly modified parameter set, where *g_IE_* =1 mS/cm^2^, such that *V_GABA_* becomes the bifurcation parameter. The other parameters remained the same. We have changed the initial parameter set to find the region of the parameter space where *V_GABA_* could play the crucial role for oscillations. We found that when *V_GABA_* is higher than –59mV, it leads to ictal oscillations (Fig. 4F). If *V_GABA_* drops below –48mV, the oscillations stop. Thus, there is substantial range of *V_GABA_* where its increase leads to the development of seizures, which might take place due to chloride accumulation before an ictal discharge (Lillis et al., 2012; Huberfeld et al., 2007).

In summary, using bifurcation analysis, we characterized the parameter regions of the model where seizure oscillations could take place. We found that transitions from seizure to rest and from rest to seizure take place via supercritical and subcritical Andronov-Hopf bifurcations. In all studied cases we found that resting and oscillatory solutions exist for large parameter variations, implying the stability of found solutions (Prinz et al., 2004; Marder et al., 2011). We showed that variations of synaptic *g_EE_*, *g_EI_*, *g_IE_*, *g_II_* and intrinsic conductances *g_AHP_* could bring the system towards seizure and move it back to the resting state. It implies that combination of reccurent synaptic currents and spike-frequency adaptation in the excitatory population accounts for the transitions between seizure and resting states.

## Discussion

The objective of this study was to investigate the role of intrinsic excitability and inhibition as mechanisms of seizure dynamics. We constructed a novel neural mass model, consisting of interacting excitatory and inhibitory neural populations driven by external synaptic input. By comparing the model with the LFP data from human hippocampal/subicular slices, we found that it could accurately represent resting states, ictal discharges, and pre-ictal oscillations after the blockade of inhibition (Huberfeld et al., 2011). Analysis of the model showed that synaptic and intrinsic conductances play the most crucial role for transitions between resting and seizure activity. By analyzing the parameter space of the model we found the oscillatory regimes specific for the resting state and seizure dynamics, and found that transitions between these regimes take place via subcritical and supercritical Andronov-Hopf bifurcations.

Starting with the pioneering work of Wilson and Cowan (Wilson et al., 1972), neural mass models have traditionally aimed to reduce the complexity of neural dynamics towards interactions between excitation and inhibition. This approach has been validated in multiple studies for describing the large-scale brain activity patterns (Jirsa et al., 2010). Additionally, it has been shown that intrinsic properties of single neurons such as spike-frequency adaptation (Fröhlich et al., 2008) substantially change spiking patterns and thus neural dynamics (Buchin et al., 2016; Bazhenov et al., 2004; Kager et al. 2000). So far these types of interactions have not been explicitly taken into account in neural mass models.

In this work we developed a novel mass model by adding AHP currents (Buchin et al., 2010) to the excitatory population. This allowed to efficiently take into account not only seizure and resting state dynamics (Wendling et al., 2012), but also pre-ictal oscillations. In our model seizure activity takes place due to imbalance between self-excitation, adaptation and inhibition. We found that reducing the amount of inhibition to the excitatory population provokes seizure activity. Nonetheless, inhibition plays an important role in orchestrating seizures as well (Fig. 2B). We found that the complete lack of inhibition leads to the development of slow oscillations with significantly different frequency content than seizures (Fig. 2C). Thus, we propose that inhibition, together with single neuron intrinsic properties provided by adaptation, plays an important role controlling the seizure dynamics.

We have investigated multiple mechanisms responsible for generation of seizure activity. In the proposed model, seizure oscillations could be generated by increased recurrent excitation *g_EE_*, decreased excitation of the inhibitory population *g_IE_*, decreased inhibition of the excitatory population *g_IE_*, increased recurrent inhibition in the inhibitory population *g_II_*. Changes in the intrinsic excitability of the excitatory population such as decrease of intrinsic adaptation *g_AHP_* and increase of the GABA_A_ reversal potential *V_GABA_* could also lead to seizure oscillations. We speculate that various physiological parameters combinations could lead to seizure activity, as found by Jirsa et al. (2014). The combination of multiple factors such as increased chloride concentration in the pyramidal cells and GABA_A_ reversal (Buchin et al., 2016; Huberfeld et al., 2011; Lillis et al., 2013), together with an increase in extracellular potassium concentrations (Bazhenov et al., 2004; Krishnan et al., 2011) and decreased activity of interneurons (Ziburkus et al., 2006), all contribute to seizure initiation. Combination of these factors and their relative contribution should be evaluated via further experiments and modeling.

Adaptation on the single neuron level could be achieved by calcium-dependent potassium currents (Bazhenov et al., 2004, Jung et al. 2001). In our model, AHP is the key mechanism for termination of population bursts during seizure oscillations (Fig. 2B) and pre-ictal discharges (Fig. 2C). The alternative potential mechanism of termination of these bursts are GABA_B_ currents provided by the inhibitory population (de la Prida et al. 2006). We predict that in the complete absence of the inhibitory neurotransmission including GABA_A_ and GABA_B_ synapses, the purely excitatory network in the epileptogenic slice of human subiculum would be capable of generating self-sustained pre-ictal oscillations due to negative feedback provided by AHP (Ratnadurai-Giridharan et al., 2014) and other intrinsic adaptation currents. Therefore the downregulation of excitatory neuronal adaptation currents such as AHP and/or functionally similar muscarinic-sensitive potassium currents (Stiefel et al., 2008) could lead to seizure initiation. According to the model the pharmacological strategy aiming to increase the amount of adaptation in the excitatory population would lead to the decreased susceptibility towards seizures.

GABA_B_ inhibition could also participate for the termination of population bursts. As shown by (de la Prida et al. 2006), the joint blockade of GABA_A_ receptors by PTX and GABA_B_ receptors by CGP led to generation of all-or-none population bursts in CA3 mouse hippocampal slices. In our experiments we did not test for the possibility that GABA_B_ could participate for the pre-ictal discharge termination. Further experiments are needed to divide the contributions of GABA_B_ and AHP for the burst termination.

Note that oscillations in the slice switched from ictal discharges to pre-ictal ones after full GABA_A_ blockage. This transition was possible only if seizures were already established in the slice (Huberfeld et al., 2011). It implies that there are excitability and synaptic plasticity changes in the slice associated with seizures before the pre-ictal discharges could be established using complete GABA_A_ blockade. When GABA_A_ blockers were applied before first seizure being generated, the pre-ictal and ictal oscillations were not established (Huberfeld et al., 2011).

Our model has several limitations compared to existing approaches (Wendling et al., 2012; Jirsa et al., 2014; Molaee-Ardekani et al., 2010). First, it is unable to describe the pre-ictal discharges taking place before seizure. The work of (Buchin et al., 2016) proposes a network explanation of pre-ictal discharges that take place before seizure transition (Huberfeld et al., 2011). To describe this activity, it was necessary to take into account the heterogeneity in the excitatory population caused by depolarizing GABA, while in the current model we did not take it into account. Therefore, pre-ictal discharges in our model could be generated only in the absence of inhibitory population. Second, particular features such as high frequency oscillations (Engel et al., 2009) relevant for seizure initiation (Quilichini et al., 2012) are not captured in our model. We speculate that this property could be taken into account by incorporating fast somatic and slow dendritic inhibition (Wendling et al., 2012). Third, our model is also unable to describe the interictal discharges, which have been explained in the other population models (Wendling et al., 2012; Chizhov et al., 2017). It has been found that interictal discharges in human subiculum require initial interneuron activation. Since in our model we impose the background synaptic input onto the excitatory population, the pyramidal cells are always activated before interneurons. It has been recently proposed in (Chizhov et al., 2017) that the interneuron population should receive background synaptic input, which would allow the reproduction of interictal discharges in neural mass models.

Pre-ictal discharges are generated before seizure initiation and in the absence of inhibition when seizures have been established in the slice (Huberfeld et al., 2011). These oscillations are still generated in the absence of inhibitory population (Fig. 2C). Using the model, we show that in this case the background synaptic input to the excitatory population *I_E_*(*t*) is necessary to generate the periodic pre-ictal oscillations. When *I_E_*(*t*) is absent, there are no pre-ictal discharges in the model, Fig. 3C. We speculate that before seizure initiation the interneurons are becoming non-functional because of depolarization block (Ziburkus et al., 2006) and GABA_A_ reversal (Lillis et al., 2013) thus allowing the pre-ictal discharges to be generated before seizure initiation (Huberfeld et al., 2011). The proposed model could explain the presence of pre-ictal discharges only in the complete absence of inhibition, Fig. 2C. The possibility of pre-ictal discharge generation before seizure due to non-functional inhibition could be investigated in the future studies.

During seizures or ictal discharges, the frequency content of spiking activity might substantially change (Fig. 3E). This can be explained using the current model as due to the gradual increase of recurrent excitation *g_EE_* (Fig. 3A) or the increase of recurrent inhibition *g_II_*. Note that the frequency content of seizure oscillations in the end of it might be similar to the pre-ictal discharges (Fig. 2C.). However, pre-ictal oscillations are possible in the model only in the absence of inhibition (Fig. 2B), as in the experimental data when the GABA_A_ synaptic activity is completely blocked.

The primary advantage of our model compared to more abstract ones such as Jirsa et al. (2014) is that it provides more firm biophysical explanations linking single neuron properties to population dynamics (Gerstner et al., 2002; Johannesma, 1968; Chizhov et al., 2007). Our approach could be extended to take into account the shunting effect of inhibition by adjusting the firing rate transfer function (Chizhov et al., 2013). To describe the additional mechanisms of seizure transition, the present model could include slow activity-dependent parameter changes similar to (Proix et al., 2014, Bartolomei et al., 2014; Ullah et al., 2009; Cressman et al., 2009). There are multiple biophysical mechanisms that could play the role of slow variable bringing the network towards seizure (Naze et al., 2015), including dynamic ion concentration of extracellular potassium (Bazhenov et al., 2004), intracellular chloride (Buchin et al., 2016, Jedlicka et al., 2011), and intracellular sodium (Krishnan et al., 2011, Karus et al., 2015) in pyramidal cells. The population model could be further modified to incorporate these slow mechanisms to describe seizure initiation.

A common problem with neural mass models in general is their limited ability to generate the experimentally measurable signals (Lytton, 2009). In this work we used the average voltage of the excitatory neural population as the approximation of the LFP signal near the neurons’ somas (Ratnadurai-Giridharan et al., 2014). Given the distant-dependence of the LFP signal, this model should be considered only as a first approximation (Buzsáki et al., 2012). More detailed approaches describing populations of two-compartmental neurons (Chizhov, 2013, Chizhov et al., 2015) could also provide better approximation for the LFP.

Epilepsy is a complex phenomenon involving the dynamic interactions between multiple components of the nervous system (Lytton, 2008,Bartolomei et al., 2014). In this work we have investigated the particular role of inhibition and adaptation and their implications for seizure dynamics. Reconciling modeling results with experimental data, we have shown that seizure activity cannot be generated in the complete absence of the inhibitory population and adaption in the excitatory population. Further development of theoretical and experimental approaches in epilepsy research may lead to a better understanding of its mechanisms and the development of new therapeutic targets.

Author contributions
Anatoly Buchin: designed research, performed research, analyzed data, wrote the paper
Cliff C. Kerr: designed research, analyzed data, wrote the paper
Gilles Huberfeld: designed research, performed research, wrote the paper
Richard Miles: designed research, wrote the paper
Boris Gutkin: designed research, wrote the paper

## Acknowledgments

We would like to thank Anton Chizhov for useful comments and criticism of our work. Initial ideas were developed during the Advanced Course in Computational Neuroscience in Będlevo, Poland (http://www.neuroinf.pl/accn).

